# αB-crystallin inhibits amyloidogenesis by disassembling aggregation nuclei

**DOI:** 10.1101/300541

**Authors:** Olga Tkachenko, Justin L.P. Benesch, Andrew J. Baldwin

## Abstract

Amyloid formation is implicated in a range of neurodegenerative conditions including Alzheimer’s and Parkinson’s diseases. The small heat-shock protein αB-crystallin (αBC) is associated with both, and directly inhibits amyloid formation *in vitro* and its toxicity in cells. Studying the mechanism of aggregation inhibition is challenging owing to sample heterogeneity and the dynamic nature of the process. Here, by means of NMR spectroscopy and chemical kinetics, we establish the mechanism by which the protein α-lactalbumin aggregates and forms amyloid, and how this is inhibited by αBC. In particular, we characterise the lifetime of the unstable aggregation nucleus, and determine that this species is specifically destabilised by αBC. This mechanism allows the chaperone to delay the onset of aggregation, although it is overwhelmed on longer timescales. The methodology we present provides a mechanistic understanding of how αBC reduces the toxicity of amyloids, and is widely applicable to other complex mixtures.

## Introduction

Neurodegenerative conditions such as Parkinson’s and Alzheimer’s diseases (PD and AD) are some of the greatest health challenges for the ageing world population^3^. These conditions are associated with aggregation of protein in the form of amyloid, which can be highly toxic^4,5^. Cells possess a sophisticated defence network against such protein aggregates that includes the small heat-shock proteins (sHSP)^6–9^. In humans, the sHSP αB-crystallin (αBC) is linked to AD and PD^10^, being up-regulated in neurons and glia adjacent to amyloids in humans^11–13^, and found within plaques analysed *ex vivo*^12,14^. *In vitro*, αBC significantly inhibits amyloid formation of a wide range of proteins^10^. The molecular mechanism of this process, comprising a description of the kinetic and thermodynamic relationships between interacting species, has proven highly challenging to obtain owing to the heterogeneity and kinetic instability of complexes formed between protein aggregates and chaperones.

Amyloid formation typically begins from monomeric protein, culminating in high order aggregates. In principle, chaperones could associate with any, or all, of these species along the amyloidogenesis pathway. A wide range of biophysical techniques including light scattering, changes in fluorescence of histological dyes, circular dichroism, size exclusion chromatography, ultracentrifugation, native mass spectrometry, small-angle X-ray scattering, and nuclear magnetic resonance (NMR) spectroscopy have been employed to determine features of this process^5,10,15–25^. Significant insights into how proteins aggregate, and how chaperones attenuate aggregation, have been obtained by relating such data to analytical equations in specific regimes^26–28^. αBC is a particularly challenging chaperone to study, as it spontaneously assembles into a polydisperse ensemble of exchanging oligomers^29–31^ (Fig. 1). If the N- and C-terminal regions are removed, the excised core domain (cαBC) forms dimers that inhibit amyloid formation similarly to the full-length, oligomeric form of the protein^1^. Despite extensive research efforts, the mechanism by which these forms of αBC interact with substrate to inhibit amyloid formation has remained elusive.

**Figure 1.**
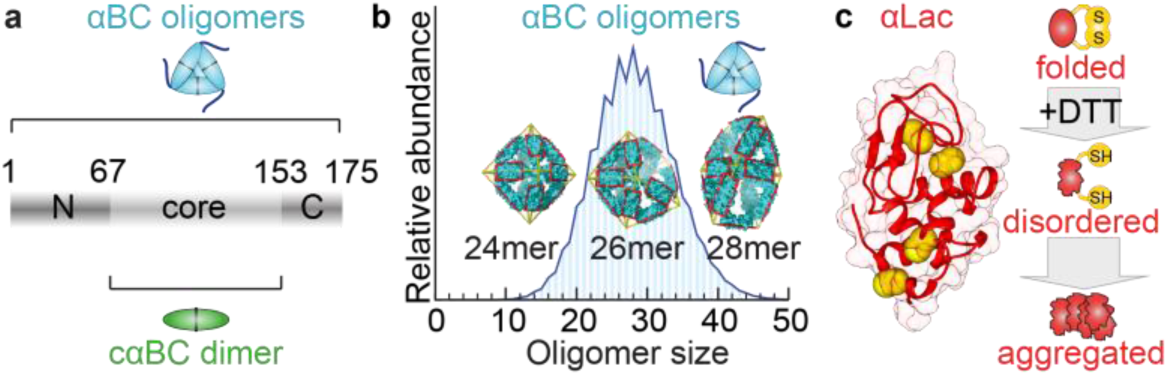
| The chaperone αBC and the aggregating client αLac. **a** The core domain of αBC is flanked by disordered N- and C-termini. The excised core exists as a dimer (cαBC). Both full-length αBC and cαBC are potent chaperones^1^. **b** Full-length αBC exists as a polydisperse ensemble and structural models are available for the principally populated oligomers^2^. **c** Structure of apo αLac with the four disulphide bonds highlighted in yellow. Addition of DTT leads to reduction of the disulphides and adoption of a disordered conformation, which, under the conditions tested here, aggregates to form amyloid.

Here, we present a novel strategy for mechanism determination based on combining NMR spectroscopy with numerical chemical kinetic modelling and stepwise regression. With differential isotope labelling, our NMR experiments allow us to monitor changes in both the chaperone and the amyloid-forming protein directly and independently. By analysing datasets acquired over a range of concentrations, we can test a variety of possible numerical aggregation models, to find those that best fit the data. Through quantitative study of the effects of αBC on a model amyloid-forming protein, bovine α-lactalbumin (αLac, **Fig. 1c**)^16^, with our method, we determine a mechanism for how a model amyloid-forming system aggregates, and how this is inhibited by αBC.

Our analysis reveals that αLac aggregates via two parallel nucleated pathways, operating on different timescales. We find that, in addition to binding mature aggregates, as has been observed previously^20, 21, 32^, the action of αBC is dominated by disrupting lowly populated aggregation nuclei, thus redirecting the substrate towards the monomeric form and thereby slowing aggregation. While both αBC and cαBC display a similar mechanism, we find that the full-length protein interacts only with aggregated forms of αLac, while cαBC additionally interacts with free monomers. This interaction with unaggregated monomers would be undesirable in the context of a cell and provides a rationale for why αBC forms oligomers. Remarkably, while both forms of αBC attenuate aggregation in a concentration-dependent manner, we find that at longer times both proteins are effectively overwhelmed, and that there a limited window of time and client concentration in which αBC is effective. This provides quantitative insight into how αBC, and potentially other sHSPs, may be involved in neurodegenerative disorders, including AD and PD. Our combined NMR and chemical kinetics methodology is applicable to the mechanistic study of other complex and heterogeneous mixtures, and provides a quantitative foundation for therapy design.

## Results

### A quantitative NMR assay for monitoring protein aggregation

To obtain a reference data set, we performed a static light scattering assay, a widely used *in vitro* measure of sHSP chaperone activity^10^. Consistent with previous data^1,16,33^, upon addition of the reductant dithiothreitol (DTT) to αLac, a sigmoidal increase in scattering was observed (**Fig. 2a**). The resulting aggregates were found to be amyloid (**Fig. S1**). Flexible fibrillar aggregates with β-sheet secondary structure were observed in electron microscopy (EM) images (**Fig. 2e**) that bound the histological dye thioflavin T (ThT) (**Fig. S1**). Addition of an equimolar amount of either full-length oligomeric αBC or cαBC resulted in complete abrogation of the increase in scattering over the timescale of the experiment (Fig. 2a). These results clearly demonstrate the potency of the chaperone in inhibiting amyloid formation.

**Figure 2.**
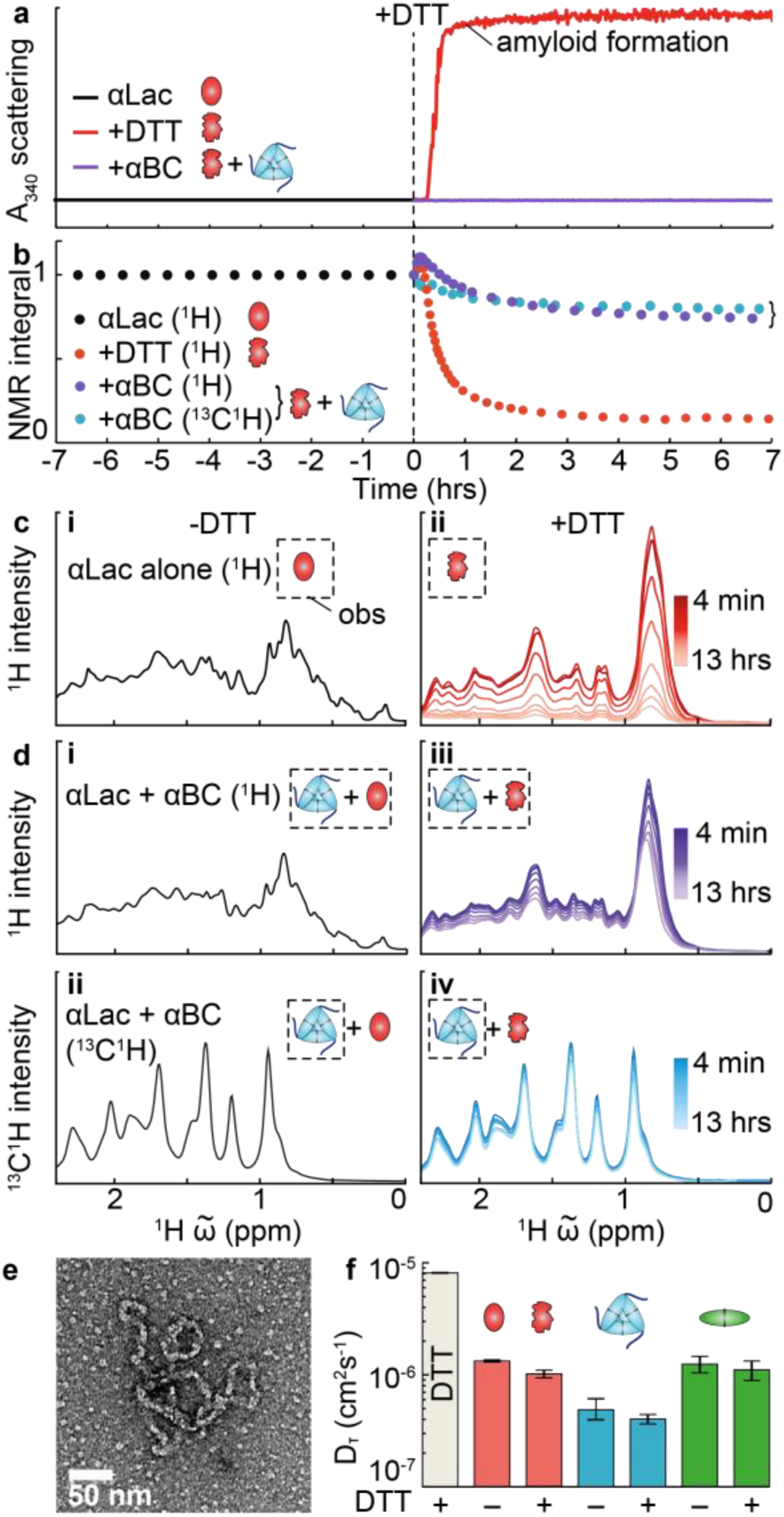
| Following amyloid formation of αLac by light scattering, and NMR. **a** Aggregation of 200 μM αLac followed by light scattering in the absence (red) and presence (purple) of αBC at a 1:1 molar ratio, **b** The same process followed by NMR, where the ^13^C isotopically enriched chaperone can be monitored independently, **c i** ^1^H NMR spectra of αLac prior to addition of DTT in the absence of chaperones. The observed species (obs) is illustrated in a dashed box. **ii** Time evolution of the αLac ^1^H spectra following addition of DTT. **d** ^1^H (**i**) and ^13^C-edited ^1^H spectra (**ii**) of the αLac/αBC mixture prior to the addition of DTT. Both species are observed in **i**, whereas only chaperone is observed in **ii**. The evolution of both spectra following DTT addition is shown in **iii** (^1^H) and **iv** (^13^C-edited ^1^H). The time-course of the integrated signal from these spectra is shown in **b. e** Negative stain electron microscopy showing aggregates of αLac in the absence of αBC following 12 hours of exposure to DTT. **f** Translational diffusion coefficients of the species observed measured by NMR. The values obtained are consistent with the spectra being dominated by the free, unbound species being observed at all times (see also **Fig. S3**).

Light scattering has a complex dependence on the shape, size, and distribution of the aggregating particles, making it challenging to analyse such data quantitatively to obtain a molecular mechanism. By contrast, NMR spectroscopy is amenable to quantitative studies of amyloid formation, as the observed signal can be directly related to the concentration of observed particles^22, 34^. In the absence of DTT, the ^1^H NMR spectrum of αLac (200 μM) is typical of a folded, globular protein^35^ (**Fig. 2c, i**). Upon the addition of DTT, the spectrum changes in appearance and returns a ~20% decrease in the translational diffusion coefficient (**Fig. 2c ii; f**). At this αLac concentration, the conformational change results in an increase in overall signal intensity to a maximum within minutes, within the apparent lag phase of the light scattering measurements. These observations are consistent with the formation of a disordered “molten globule” ^36–39^. The signal then decays on a timescale of hours to a steady-state value. This is consistent with expectations that high molecular mass aggregates are formed over time that are not directly observable by solution NMR owing to their relatively rapid transverse relaxation^16, 40^. We found the data to be very reproducible at a range of αLac concentrations between 100-2000 μM (**Fig. S2**). At higher concentrations, the initial rise in signal intensity was attenuated, the signal decay rate increased, and the steady-state relative abundance of monomer decreased (**Fig. S2**). The total αLac concentration dependence of residual monomer levels at the end of the experiment demonstrates that αLac aggregation must be a reversible process, or else the signal ought to decay to zero as all monomers are converted into aggregates^23^. These results demonstrate that the NMR experiment provides a simple assay to follow the change in free monomer concentration quantitatively and in real time.

### Selective isotope labelling allows independent monitoring of chaperone activity

To extend our approach to a multi-component mixture, we employed a selective isotope-labelling strategy so that, by performing isotope-edited NMR experiments, we would be able to monitor the components independently. For full-length αBC, we introduced ^13^C labels, allowing us to specifically follow the flexible C-terminal region on the exterior of the oligomers^41–43^. For cαBC, we employed ^15^N enrichment, allowing us to follow each position in the backbone of the protein. These were mixed individually with unlabelled αLac. We could therefore monitor both the chaperone and amyloid formation independently, without the need for extrinsic labels (**Fig. 2b and d, Fig. S2**). Prior to addition of DTT, the ^1^H spectra of the mixtures were the sum of the spectra of the individual components (**Fig. S3**). After the addition of DTT, even at the earliest times, where the light scattering traces in the presence or absence of chaperone are indistinguishable (and approximately zero, Fig. 2a), significant changes were seen in the NMR spectra, revealing that the experiment is sensitive to early events in aggregation (**Fig. S2**). In addition, the rate of αLac signal decay was substantially slower in the presence of chaperone, indicative of chaperone activity (**Fig. 2b**).

No significant change was observed in either the position or linewidth of resonances in two-dimensional isotope-edited NMR spectra of either chaperone (**Fig. S3**). Instead, a uniform loss of signal intensity was observed across the spectrum (**Fig. 2d, Fig. S3**) indicating that the observed species are in the ‘slow’ regime of chemical exchange^34^. Similarly to αLac alone, diffusion experiments revealed that the observed species were free in solution, rather than trapped in aggregates (**Fig. 2f** and **S3**), and that the loss in signal intensity coincided with the build-up of high molecular-mass aggregates visible in EM images (**Fig. S1**).

We acquired a large dataset of interleaved NMR experiments by varying αLac concentration systematically (**Fig. S2**). Both ^1^H and isotope-edited NMR signal intensities reached non-zero, concentration-dependent steady-state values at longer times, revealing the reversible nature of chaperone-αLac interactions (**Fig. S2**). Significantly more signal was observed at longer times after incubation with chaperone, indicating enhanced protection (**Fig. S2**). The visible differences in the data effected by the addition, and variation, of chaperone reveal that the NMR experiment is information-rich, and sensitive to molecular processes involved in aggregation.

### Chemical kinetics modelling reveals that αLac aggregates via two parallel nucleated pathways

To elucidate the quantitative mechanisms of amyloid formation and chaperone action from our data, we chose to develop a chemical kinetics modelling strategy combined with stepwise regression. In our approach, specific aggregation mechanisms of varying complexity are hypothesised, and simulated (**Table S2**). Each proposed mechanism predicts how the concentration of species of every stoichiometry varies with time for a given set of rate constants, allowing us to simulate the NMR timecourse data. The rate constants for a model are then optimised numerically via a multi-step stimulated annealing protocol to determine best-fit values and the lowest possible χ^2^ to the global dataset that the model could achieve. A combination of forward (increasing complexity) and reverse (reducing complexity) steps were used to compare and construct models, where complexity refers to the number of fitting parameters (**Fig. S4**). To avoid the inherent risks of over-fitting to which stepwise regression is prone, F-tests were performed to determine whether any improvement in fit with an increased number of parameters could be explained by noise in our data (P-level < 0.05). In addition, to test the resulting mechanisms, a subset of data was deliberately excluded from model building and was subsequently simulated using the parameters obtained from the main set of data.

We first employed this approach to investigate the aggregation of αLac in the absence of chaperone. The simplest model for amyloid formation is linear polymerisation, which assumes that aggregation proceeds through consecutive monomer addition, and that all association and dissociation rates of oligomers are independent of stoichiometry^44^. This model, parameterised by two rate constants, provided a poor description of the data, and could be confidently excluded (**Fig 3a, Table S2**, model 1). We then built additional complexity into the model, including effects such as amyloid formation proceeding via an aggregation-prone monomer form, fragmentation of the aggregates, variations in the size of the aggregation nucleus, spherical rather than linear aggregate growth, and multiple aggregation pathways (**Table S2**). Overall, we tested 17 mechanisms of varying complexity until we identified a model that explained our data well, and where increasing the model complexity further did not result in a statistical improvement.

**Figure 3.**
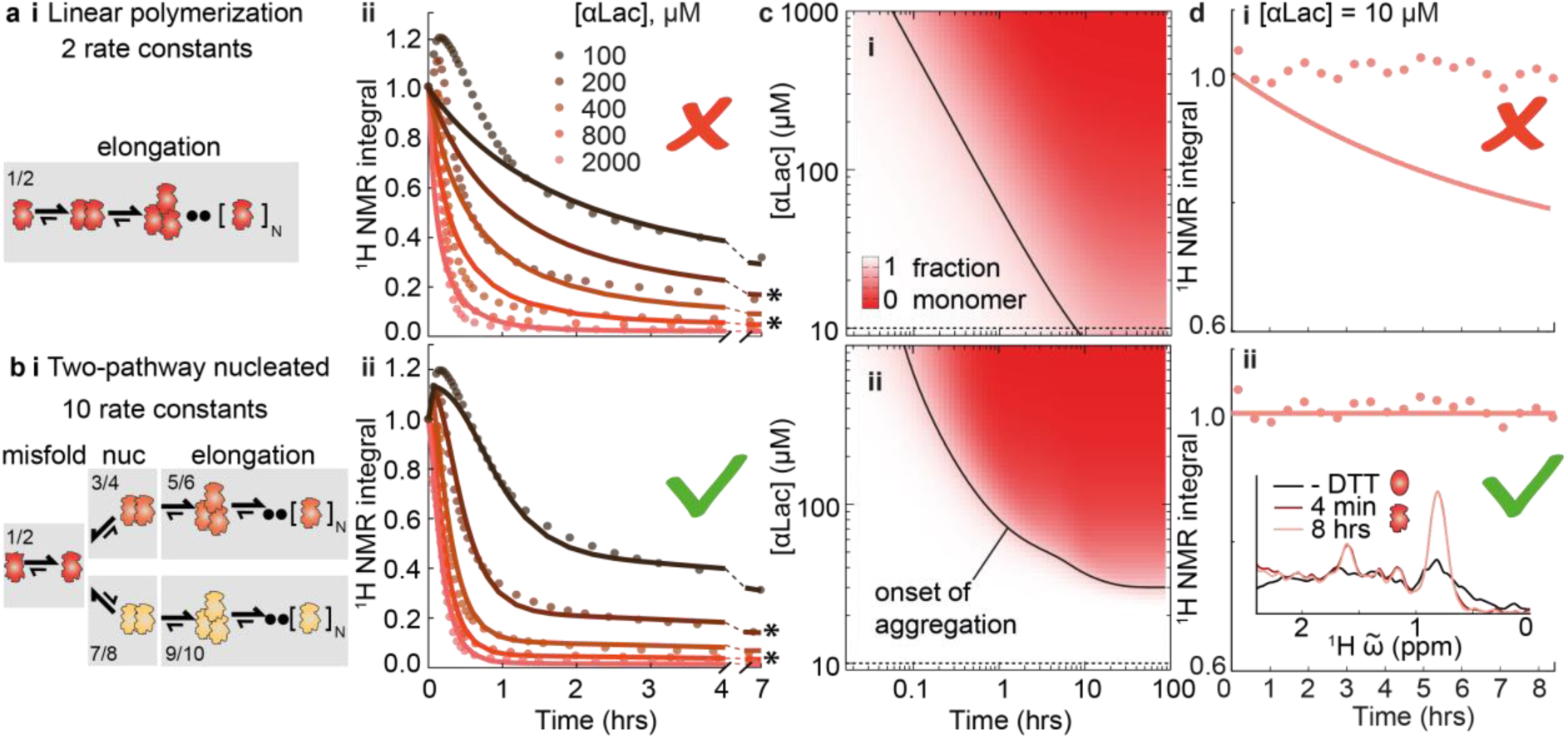
| Determining the mechanism of amyloid formation by αLac. **a i** The simplest aggregation model, linear polymerisation, parameterised by two rate constants. **ii** Aggregation timecourses at five αLac concentrations (circles) with the fitted linear model (lines), revealing a poor fit to the data. **b i** The optimal aggregation model determined by our analysis (**Table S2**), parameterised by 10 rate constants. **ii** The fit of the optimal model to the data. Concentrations marked with an asterisk (200 and 800 μM) were excluded from the analysis and simulated only using the final rate constants for validation purposes. **c** An aggregation surface describing the loss of free monomer as a function of time and total protein concentration for (**i**) the linear polymerization model and (**ii**) the optimal model. The 80% free monomer isochor indicating the onset of aggregation is plotted (black line). The expected timecourse at 10 μM αLac is indicated as a line. **d** The experimentally measured timecourse at 10 μM αLac (circles) and the predictions for (**i**) the linear polymerization model and (**ii**) the optimal model. The inset in **ii** shows an overlay of representative NMR spectra for this dataset, illustrating the absence of signal loss over time in accordance with our optimal model.

Achieving a satisfactory global fit was remarkably challenging (**Fig. 3a, Table S2**), with low complexity models unable to explain the data, revealing the high information content of the NMR data. A model of intermediate complexity, characterised by 10 rate constants was identified as our optimum in terms of lowered χ^2^ values versus the increase in the number of model parameters. This model includes formation of an on-pathway aggregation-prone monomeric species, and two parallel nucleated aggregation pathways (**Table S2, Fig. S4, S6**). Each of these aspects can be related directly to features in the raw data: without including an aggregation-prone monomer, we could not account for the increase in signal at early times; without nucleation we could not properly account for the concentration dependence of signal decay rate at early times; and without two pathways we could not account for the bi-phasic decay of signal at long times (**Fig. 3b**). The interrogation of the NMR aggregation curves using chemical kinetics therefore provides an in-depth, quantitative understanding of the αLac aggregation mechanism, identifying and characterising key species (**Fig. 3b**).

### αBC inhibits αLac aggregation by disassembling aggregation nuclei

Having determined the mechanism of αLac aggregation, we sought to investigate how αBC effects inhibition of this process by applying our chemical kinetics approach to the data in the presence of chaperone. We performed extensive model testing to find one that could globally explain our data for each chaperone (**Fig. 4, Fig. S4, Tables S3-5**), simulating both the chaperone and aggregate data ‘channels’. The parameters for αLac aggregation were fixed to those derived in the analysis of αLac alone (**Fig. 3, Fig. S6**). We then introduced additional steps involving interactions between chaperone and the various aggregate states of αLac. 40 different association models parameterised by 2 to 12 rate constants were tested to determine if they can fit to the data (**Tables S3-5**). As with αLac, forward selection was performed, where a series of models of increasing complexity were tested. This process identified complex models that produced good fits to the data. These models were then subjected to reductive elimination, where individual binding processes were excluded, to obtain a minimally complex model that explained the data adequately.

**Figure 4.**
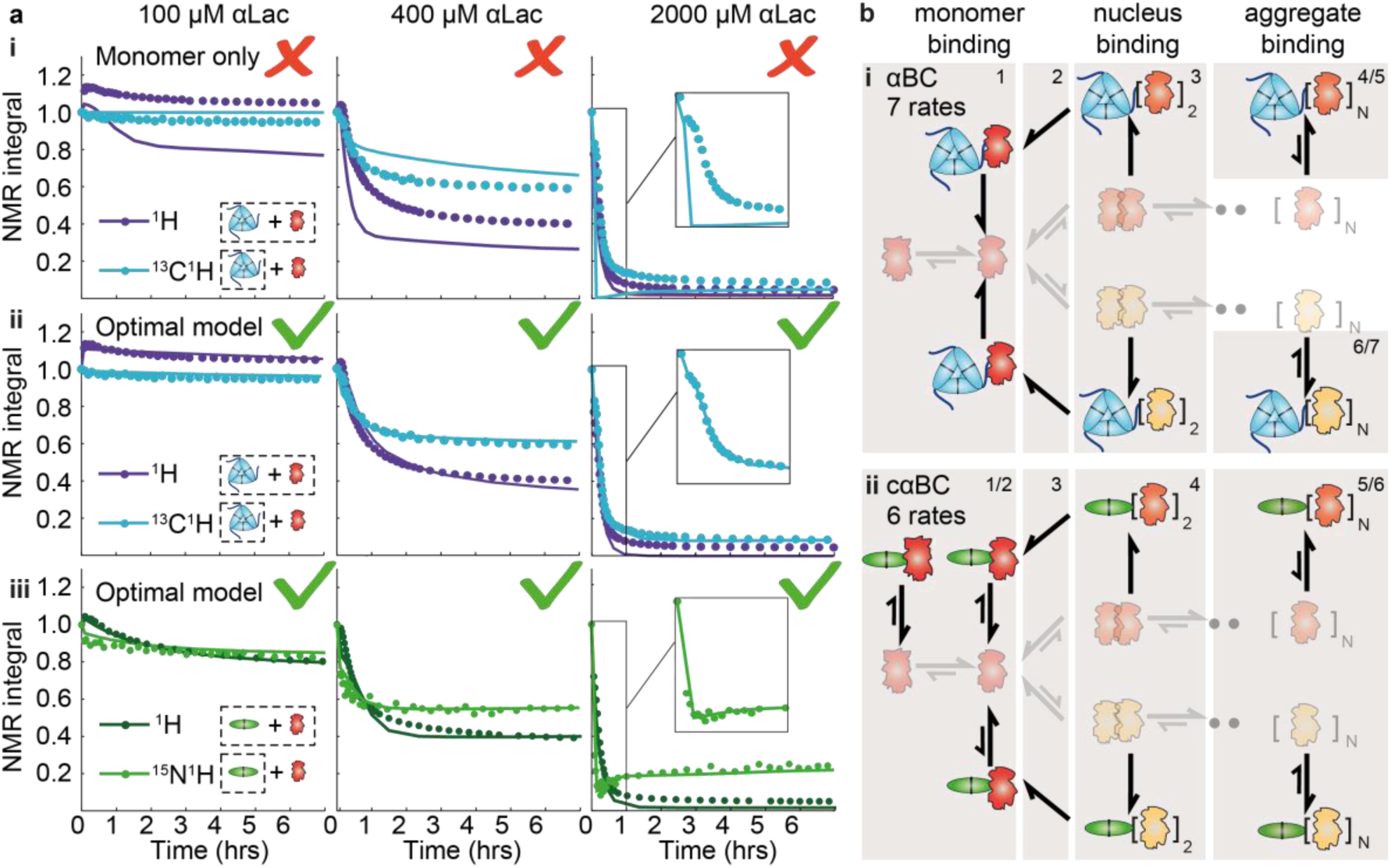
| Mechanism of αLac amyloid inhibition by αBC and cαBC. **a** NMR signal intensity timecourses for three concentrations of αLac in the absence and presence of either αBC or cαBC at 200 μM. For each αLac concentration, isotope-unedited ^1^H NMR signal and ^15^N-(cαBC) or ^13^C-(αBC) edited chaperone-only ^1^H signal is shown as circles. Corresponding lines show fits from (**i**) a model where the chaperone can bind only monomeric αLac, (**ii**) and (**iii**) optimal models for αBC and cαBC respectively. **b** The corresponding mechanisms for the optimal models (**i** – αBC, **ii** – cαBC).

We first tested the hypothesis that reversible binding of the chaperone to various species along the aggregation pathway effectively stalls the aggregation process. We tested chaperone binding to all species equally, or separately to misfolded αLac monomers, nuclei or aggregates. These simple models that allowed binding of one type of species, or increasingly complex models that included all combinations of binding effects were in poor agreement with the data (**Fig. 4a i, Table S3**). This revealed that the NMR data could not be explained only by competitive binding of the chaperone to aggregating αLac. To account for the persistence of αLac monomer signal in our data, we hypothesised that the chaperones could additionally disrupt the aggregation nuclei after binding (**Fig. 4b** and **Fig. S7**), resulting in chaperone-bound monomers, which can then dissociate. Remarkably, we found that models including this effect were in excellent agreement with our data for both αBC and cαBC, at all concentrations surveyed and throughout the reductive elimination process (**Tables S3-5, Fig 4a** and **Fig. S5**). The final models, selected through our statistical criteria, required 6 rate constants for cαBC, and 7 for αBC to explain the entire NMR dataset (**Fig. 4b, Fig. S8**). For both chaperone forms, it was necessary to include binding to both the nucleus and higher order aggregates, and a pathway for nucleus disassembly. A notable difference between the two forms of the chaperone was that cαBC bound to all types of species, whereas αBC bound only the aggregating oligomers of αLac but not the monomer, revealing it to be more selective than the truncated form.

To validate the final models, over and above the data-partitioning strategy (**Fig. S5**), we used the fitted parameters from the best-fitting models to test consistency with the form of the 2D NMR spectra, where both chaperones were observed to be in slow chemical exchange (**Fig. S3**). The fastest predicted exchange rate from our model was determined to be 0.2 s^-1^ (**Section S1.8**), which is consistent with the observed slow exchange. In addition, a notable prediction of the nucleated aggregation model of αLac is that <10 μM, no significant aggregation will occur on the timescale of days. We independently tested this prediction, and found that this was the case (**Fig. 3c-d**). Finally, EM images obtained 12 hours after the addition of DTT show two clearly distinguishable morphologies of aggregate: large filamentous structures, and smaller spherical aggregates (**Fig. 2e** and **Fig. S1c**). This is consistent with prediction of two parallel aggregation pathways for αLac (**Fig. S6**)^33, 45^.

### αBC kinetically stabilises aggregating αLac

Our final models for αBC/αLac interaction allow us to simulate the proportion of free αLac monomer expected at any given time and protein concentration, and how this is modulated by interaction with the chaperones. We defined an ‘excess monomer factor’ as the ratio of free αLac present in solution relative to that expected in the absence of chaperone (**Fig. 5**). Calculation of the excess monomer factor, over a broad range of αLac concentrations and as a function of time, reveals a bounded region that we term a ‘protection window’ (**Fig. 5a**). At early times, there is little aggregation, and hence no chaperone activity. In the protection window, both chaperones significantly delay the onset of aggregation (**Fig. S9**), leading to accumulation of additional αLac monomer and therefore rescue factors above 1 on the timescale of minutes to hours (**Fig. 5**). Most notably, at longer times, this protection is lost, as the thermodynamically driven aggregation process ultimately overwhelms the chaperone. This occurs earlier for higher αLac concentrations (**Fig. 5**). Interestingly, the chaperone is most effective when only a small fraction of it is bound at a given time, and becomes overwhelmed when it is almost fully bound (**Fig. S9b**), consistent with a kinetic effect. The protection window of αBC was found to be significantly larger than that of cαBC at the same concentration, indicating that its increased selectivity and more fine-tuned association and dissociation rates (**Fig. S8**) make it ultimately a more effective chaperone. In terms of mechanism, nucleus disassembly and aggregate binding were the key chaperone actions giving rise to the protection window (**Fig. S10**).

**Figure 5.**
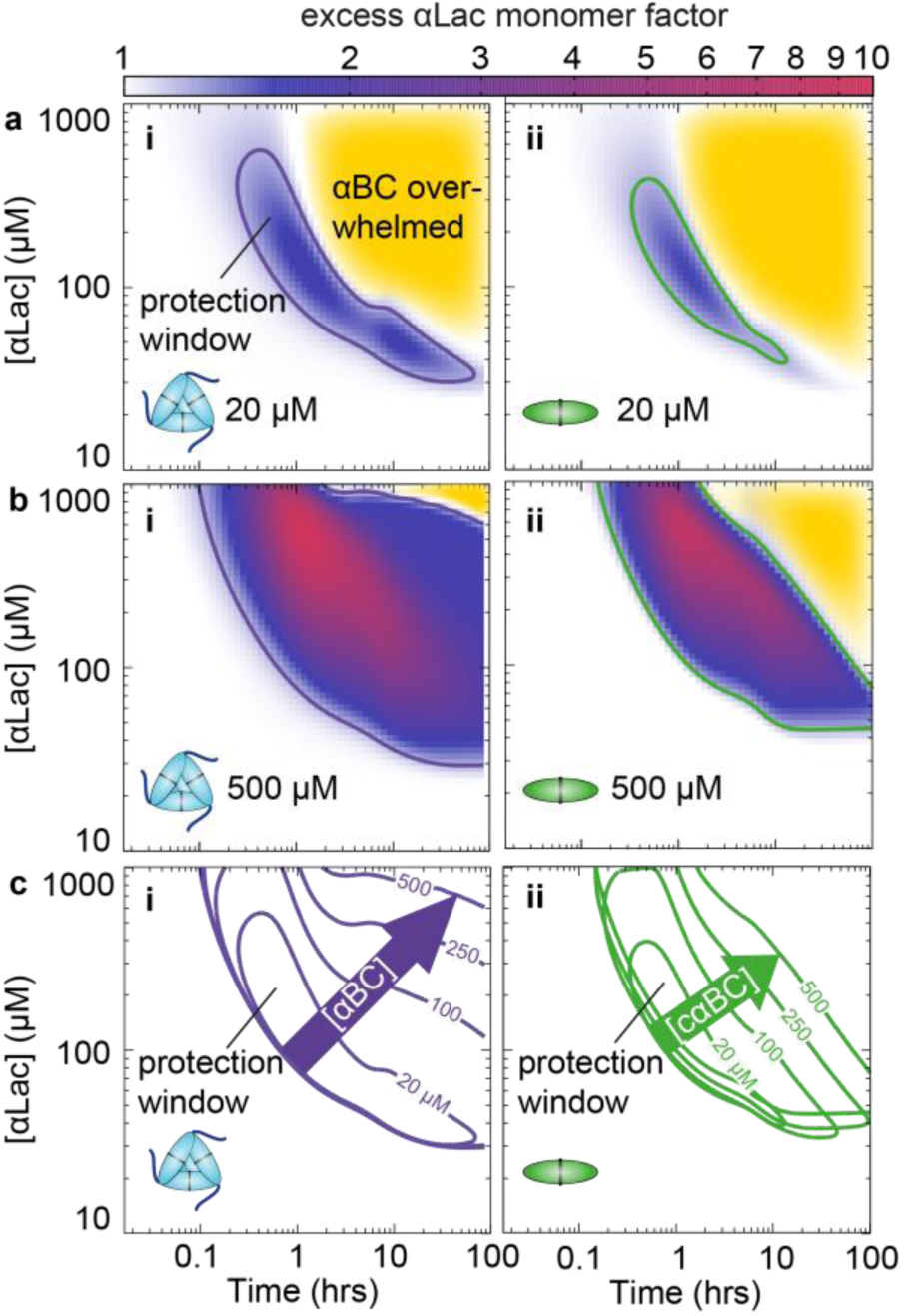
| Effect of αBC (i) and cαBc (ii) on αLac monomer. Protection is expressed as the fold excess αLac monomer, versus total concentration of αLac and time for 20 μM (**a**) and 500 μM (**b**) chaperone. In each case, a window of monomer protection is revealed, and at longer times, the thermodynamically favoured aggregates dominate (yellow). Although their action is predominantly kinetic, both chaperones attenuate the point of onset of aggregation significantly (see also **Fig. S9**). Contour lines are drawn at the value of 1.2 (20% increase in free αLac monomer). **c** The 20% increase contour line shown as a function of chaperone concentration, illustrating the window of αLac monomer protection by the chaperones.

## Discussion

The combined NMR and chemical kinetics methodology we have presented here provides a quantitative means for mechanistic characterisation of protein aggregation, and its inhibition. The overall form of the real-time data varies significantly with protein concentration, which lends the approach high discriminatory power necessary for mechanism determination. In addition, NMR spectra are sensitive to changes in monomer concentration at early times, during the lag phase of traditional assays based on light scattering, demonstrating that our approach is particularly suited to the analysis of early stages in protein aggregation, which is typically the period where toxic early-aggregates are expected to form^5^. We capitalised on these capabilities to determine the mechanism by which αLac forms amyloid, and how this is attenuated by both oligomeric αBC and cαBC (**Fig. 6**). We tested a total of 57 models, requiring >1500 hours of CPU time, in order to find optimal fits to the data. These final models should be considered minimally complex, in the sense that we can unambiguously reject simpler models that lead to poor descriptions of the data, and on the other hand our data do not statistically justify the inclusion of further complexity. The final models therefore represent the most appropriate mechanisms, and crucially provide quantitative chemical explanations (in terms of rate constants) for the key features observed in the raw data.

**Figure 6.**
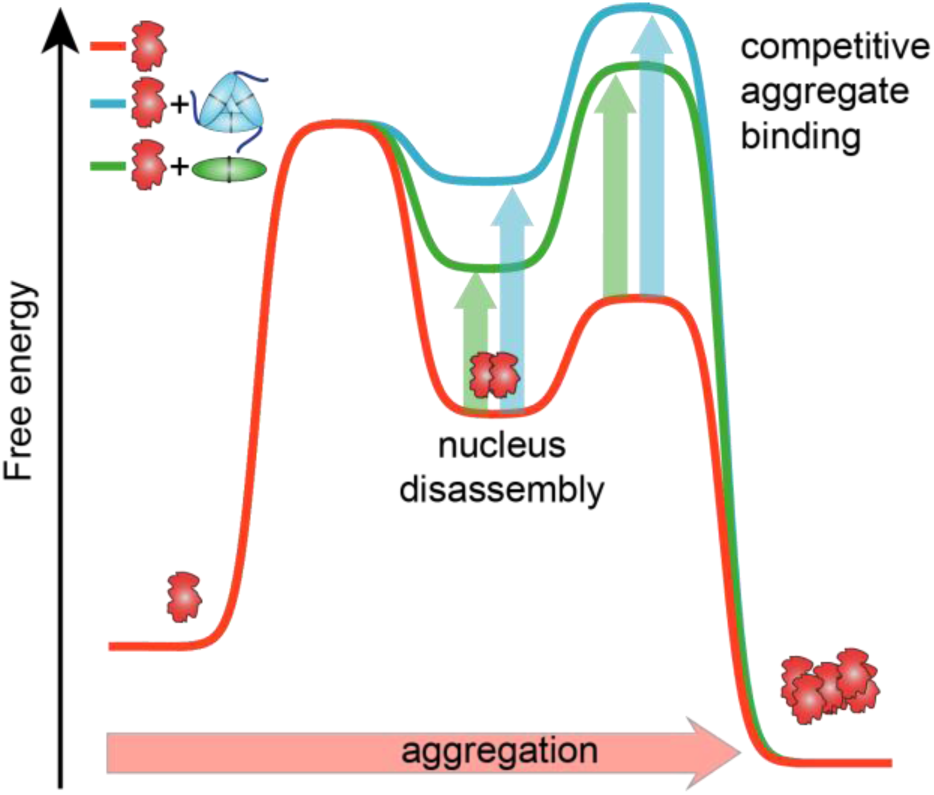
| An illustrative summary of the aggregation inhibition mechanism of αBC (blue) and cαBC (green). As the concentrations of the relevant species vary over the course of the aggregation reaction, equilibrium free energy diagrams can only approximate the chemical changes. The kinetic barrier in αLac aggregation is the formation of primary nuclei. Both chaperones effectively destabilise the primary nuclei by disassembling them, thus significantly slowing down the onset of aggregation. The chaperones also bind higher order aggregates to further slow their elongation. At long times, however, the thermodynamically favoured aggregation process eventually overwhelms the chaperones.

Our analysis provides a mechanism for αLac aggregation, where initial unfolding of the molten globule^16^ to an unstable aggregation prone monomer is followed by formation of unstable aggregation nuclei with ca. 0.2-1.2 mM dissociation constants and association rate constants on the order of 200-700 M^-1^ s^-1^ (**Fig. S6**). A consequence of this instability is that aggregation at low concentrations is very slow, but, once formed, the nuclei grow by elongation into aggregates with two distinct morphologies (**Fig. S1**), which has been observed previously ^33, 45^. The mechanism is reversible^23^ with a steady-state population of free, monomeric αLac remaining even at long times. Non-equilibrium phase diagrams (**Fig. 3c**) illustrate how the proportion of monomer is expected to develop as a function of αLac concentration and time.

We reveal that the chaperone has two major roles. The first, reported previously for αBC for different substrates^20, 21, 32^, is to bind larger aggregates and competitively stall their elongation. The second, more surprising observation is that αBC binds and disassembles the unstable aggregation nuclei. Most remarkably, this is accomplished without the use of ATP, and so αBC can be considered a passive disaggregase. Although examples of ATP-independent disaggregases are known^46^, and αBC has been shown to promote dissociation of apolipoprotein C-II fibrils^32^ and β2-microglobulin oligomers^24^, the specific destabilisation of nuclei is a previously unidentified function of sHSPs. The mechanism of nucleus disassembly provides an elegant means for curtailing amyloid formation, and, since lower molecular mass aggregates have been identified as more toxic than the higher molecular mass, mature aggregates^5^, this represents an efficient way for αBC to exert its protective role. The ability of the chaperones to inhibit aggregation is limited, and at long times in our assays, the chaperones are overwhelmed as the thermodynamically favoured aggregation dominates. At a given concentration, this protection window represents a period of time during which the chaperone has kinetically stabilised the amyloidogenic protein. This is in line with the function of sHSPs acting as “paramedics”, stabilising substrates for sufficient time such that ATP-dependent proteostatic mechanisms can be mobilised^47^.

In the main, sHSPs assemble into high molecular mass oligomers. Given that the truncated dimeric forms of the protein can also be potent chaperones, it is unclear why it is desirable for sHSPs to oligomerise. Here we demonstrate that the wild-type, evolutionarily optimised, oligomeric αBC binds only the nuclei and higher-order aggregates whereas the truncated construct, cαBC, in addition binds free unaggregated αLac monomers. In the cellular milieu, unproductive interactions with other cellular components would be undesirable. Overall, αBC maintains a greater proportion of the client in non-aggregated, monomeric form over time compared to cαBC, revealing it to be a more effective chaperone, and implicates involvement of the termini in chaperone activity.

In summary, our methodology has provided detailed insight into the mechanism by which αBC prevents the amyloidogenesis of αLac and revealed a new tactic used by chaperones, highlighting that cells have evolved a striking diversity of complementary mechanisms with which to target protein aggregation, both during ageing and times of acute stress. Our approach is suitable for the studies of a wide range of biomedical targets associated with aggregation and amyloid formation.

## Materials and Methods

### Protein preparation

Full-length human αB-crystallin (αBC, UniprotKB ID P02511) and the core domain construct (residues 68-153, cαBC) were prepared as described previously^25,48^ in either LB or M9 minimal media supplemented with ^15^NH_4_Cl or [U-^13^C]-glucose (Cambridge Isotope Laboratories). Bovine α-lactalbumin was purchased from Sigma Aldrich (L6010).

### Aggregation reactions

Aggregation reactions were performed at 37°C in 50 mM sodium phosphate pH 7.0, 100 mM NaCl and 2 mM EDTA with 5% D_2_O. Aggregation was initiated by addition of 20 mM DTT.

### NMR experiments

NMR experiments were performed on a narrow-bore Varian 14.1 T NMR spectrometer with a 5-mm z-axis gradient triple resonance room temperature probe for solution samples. The following parameters were typically used. ^1^H watergate: 32 scans per fid, acquisition time 100 ms, recycle delay 1 s. ^15^N^1^H 1D HSQCs: 72 scans per fid, acquisition time 67 ms, recycle delay 1 s, transmitter offset in ^15^N dimension 117 ppm. ^13^C^1^H 1D constant-time HSQCs: 64 scans per fid, acquisition time 64 ms, recycle delay 1 s, transmitter offset in ^13^C dimension 35 ppm. The duration of the experiments was 40 s for ^1^H watergate, 81 s for ^15^N^1^H HSQC and 73 s for ^13^C^1^H HSQC. Spectra were analyzed using NMRPipe^49^ and Sparky^50^. Processing included digital water suppression where necessary, exponential apodization at 10 Hz, phase correction and linear baseline correction. The following ranges were taken for integration (no buffer components present): ^1^H spectra: 0.77-2.73 ppm, ^15^N^1^H spectra: 6.5-9.5 ppm, ^13^C^1^H spectra: 0.84-2.33 ppm.

Pulsed field gradient diffusion experiments were carried out over 11 gradient strengths with 72 (^1^H experiments), 240 (^15^N^1^H experiments) or 64 (^13^C^1^H experiments) scans per transient and an acquisition time of 200 ms (^1^H), or 64 ms (^15^N^1^H or ^13^C^1^H) respectively. The maximum gradient strength was 60 G cm^-1^. The gradient duration was 2 ms (^1^H), 1.4 ms (^15^N^1^H) or 1 ms (^13^C^1^H) with a 200 ms delay between gradients. Each ppm point was analysed independently and the diffusion coefficient and its error were determined as the mean and standard deviation of a Gaussian distribution obtained from a histogram of measurements over the integrated ppm region.

### Simulations and model fitting

Complete kinetic models were constructed to follow the change in concentration of individual oligomeric species. Up to 4,000mers were individually considered with 10^5^ time steps. The models were parameterised by a number of rate constants (**Section S1.5**). The exact rate equations were integrated numerically and used to simulate the experimental data (**Section S1.5**). The free parameters were optimised using an algorithm that combines steepest descent (Levenberg-Marquardt) and simulated annealing protocols (**Section S1.5**). The *χ*^2^ values of the fits to the experimental NMR timecourses were each compared using the F-test (P level <0.05, **Section S1.7**). The errors in the parameters were estimated by performing the fit for each of the five concentrations independently (**Section S1.8**).

### Electron microscopy

Transmission electron microscopy was performed on a Tecnai 12 (FEI) transmission electron microscope operated at 120 kV. Samples at 5 μM in 5 μl were loaded on glow-discharged carbon grids, washed with water and stained with 2% uranyl acetate for 10 s. Images were acquired with a Gatan OneView CMOS camera.

## Acknowledgements

Simulations and parameter optimisation was performed at the University of Oxford Advanced Research Computing (ARC) facility (*http://dx.doi.org/10.5281/zenodo.22558*). The EM work was performed at the Dunn School of Pathology Imaging Facility and we thank the facility manager Errin Johnson for guidance. The FTIR experiments were performed by Robert Jacobs at the Surface Analysis Facility, Department of Chemistry, University of Oxford. We thank Henrik Müller for help with the analysis of diffusion data and with performing the ThT assays. We thank the EPSRC grant EP/J01835X/1 and the BBSRC grant BB/J014346/1 for funding. OT holds a Lamb and Flag Scholarship from St. John’s College, University of Oxford. JLPB holds a Royal Society University Research Fellowship, and AJB a David Phillips Fellowship from the Biotechnology and Biosciences Research Council.

